# Genetic polymorphisms do not predict inter-individual variability to cathodal transcranial direct current stimulation of the primary motor cortex

**DOI:** 10.1101/2020.06.13.150342

**Authors:** Michael Pellegrini, Maryam Zoghi, Shapour Jaberzadeh

## Abstract

High variability between individuals (i.e. inter-individual variability) in response to transcranial direct current stimulation (tDCS) has become a commonly reported issue in the tDCS literature in recent years. Inherent genetic differences between individuals has been proposed as a contributing factor to observed response variability. This study investigated whether tDCS inter-individual variability was genetically mediated. A large sample-size of sixty-one healthy males received cathodal-tDCS (c-tDCS) and sham-tDCS, of the primary motor cortex at 1mA and 10-minutes via 6×4cm active and 7×5cm return electrodes. Corticospinal excitability (CSE) was assessed via twenty-five single-pulse transcranial magnetic stimulation motor evoked potentials (MEP). Intracortical inhibition (ICI) was assessed via twenty-five 3ms inter-stimulus interval (ISI) paired-pulse MEPs, known as short-interval intracortical inhibition (SICI). Intracortical facilitation (ICF) was assessed via twenty-five 10ms ISI paired-pulse MEPs. Gene variants encoding for excitatory and inhibitory neuroreceptors were determined via saliva samples. Pre-determined thresholds and statistical cluster analyses were used to subgroup individuals. Two distinct subgroups were identified, ‘responders’ reducing CSE following c-tDCS and ‘non-responders’ showing no reduction or even increase in CSE. Differences in CSE between responders and non-responders following c-tDCS were not explained by changes in SICI or ICF. No significant relationships were reported between gene variants and inter-individual variability to c-tDCS suggesting the chosen gene variants did not influence the activity of the neuroreceptors involved in eliciting changes in CSE in responders following c-tDCS. In this largest c-tDCS study of its kind, novel insights were reported into the contribution genetic factors may play in observed inter-individual variability to c-tDCS.

## Introduction

Variability in response to transcranial direct current stimulation (tDCS) between individuals has gained popularity as an issue to the tDCS literature in recent years. Termed inter-individual variability, it defined the dichotomous nature by which individuals respond to administered tDCS, with some individuals responding as expected and others not responding as expected (see overview in ref. Horvath et al., 2014; Li et al., 2015; Pellegrini et al., 2018a, 2018b; Ridding and Ziemann, 2010). For the long-term potentiation (LTP)-like effects of excitatory anodal tDCS (a-tDCS) (Nitsche et al., 2007, 2003; Nitsche and Paulus, 2000), individuals displaying increases in overall corticospinal excitability (CSE) are categorised as ‘responders’ while those showing no increase or even reductions in CSE are categorised as ‘non-responders’. Conversely, the long-term depression (LTD)-like effects of inhibitory cathodal tDCS (c-tDCS) elicit reductions in overall CSE in ‘responders’ and no reduction or even increases in ‘non-responders’. A number of previous large-scale studies have investigated inter-individual variability to a-tDCS (Bashir et al., 2019; Chew et al., 2015; López-Alonso et al., 2015, 2014; Puri et al., 2016, 2015; Strube et al., 2016, 2015; Tremblay et al., 2016; Wiethoff et al., 2014), however only a small number have investigated the same phenomenon in c-tDCS.

To-date, just three previously published studies have investigated inter-individual variability in c-tDCS, whereby responders and non-responders were identified (Labruna et al., 2016; Strube et al., 2016; Wiethoff et al., 2014). Following similar electrode montages and stimulus parameters, comparable responder and non-responder rates were reported via two different subgrouping techniques between Wiethoff et al, (Wiethoff et al., 2014) and Strube et al, (Strube et al., 2016), while the dichotomous breakdown of the responses to c-tDCS were not reported by Labruna et al, (Labruna et al., 2016). Via statistical cluster analyses and whether or not an individual’s post-DCS normalised grand average response was below 1mV, approximately just 48% of healthy individuals responded to c-tDCS, at the commonly administered stimulus parameters, with reductions in CSE (Strube et al., 2016; Wiethoff et al., 2014). This being the case, there has not been thorough investigation into mechanisms behind why just 48% of individuals respond to c-tDCS as expected, and why 52% do not. This is of particular importance and relevance to the use of c-tDCS in the clinical setting where allocation of resources is crucial and predictability, reliability and reproducibility of therapeutic effects is essential.

Previous investigations into mechanisms behind inter-individual variability have focussed on a number of factors. Technical factors specific to tDCS protocols and stimulus parameters such as adjusting stimulus duration (Monte-Silva et al., 2013; Puri et al., 2016, 2015) or current intensity (Ammann et al., 2017; Batsikadze et al., 2013; Chew et al., 2015; Tremblay et al., 2016) have been investigated, as well as sensitivities to transcranial magnetic stimulation (TMS), the CSE assessment tool (Labruna et al., 2016). In addition to these, there has been a growing narrative in the tDCS literature that not all individuals that participate in studies are the same and have intrinsic differences that may contribute to inter-individual variability. For comprehensive reviews on intrinsic factors contributing to inter-indivdual variability, refer to (Li et al., 2015; Pellegrini et al., 2018a; Ridding and Ziemann, 2010).

Variations in genes that encode for regulators of cortical plasticity are one factor that has previously been investigated. Variations in a gene that encodes for the nerve growth factor brain-derived neurotrophic factor (BDNF) and their effects on response to non-invasive brain stimulation (NIBS) and TMS measures of CSE and cortices-cortical excitability have been extensively investigated (Antal et al., 2010; Brunoni et al., 2013; Chhabra et al., 2016; Di Lazzaro et al., 2015; Frazer et al., 2016; Fujiyama et al., 2014; Hwang et al., 2015; Puri et al., 2015; Teo et al., 2014). However with lack of definitive conclusions on the role BDNF in LTP-like and LTD-like cortical plasticity following NIBS protocols such as tDCS (Brunoni et al., 2013; Chhabra et al., 2016; Di Lazzaro et al., 2015; Fujiyama et al., 2014; Mastroeni et al., 2013) has encouraged investigations into gene variations of other regulators of cortical plasticity. The LTP-like mechanisms of a-tDCS led a recent large-scale study to investigate the role of genetic variations in the genes that encode for N-methyl-D-aspartic acid (NMDA) receptors and gamma-Aminobutyric acid (GABA) receptors and their predictive value for a-tDCS inter-individual variability (Pellegrini et al., 2020a). Novel large-scale investigations into genetic variations and their predictive value for c-tDCS inter-individual variability are yet to be conducted in the tDCS literature.

The basis for investigating inter-individual variability to c-tDCS lies in its potential use as an adjunct or alternative therapy to neurological populations such as those who experience excititoxicity such as Epilepsy and seizures as well as psychological populations such as those who experience anxiety disorders. A number of recent case reports and studies have reported improvements in seizure severity and frequency following trials of c-tDCS (Assenza et al., 2017; Lin et al., 2018; Tecchio et al., 2018; Tekturk et al., 2016; Yook et al., 2011; Zoghi et al., 2016) as well as fear response extinction in anxiety populations (Ganho-Ávila et al., 2019). With drugresistance a common problem in those who suffer seizures (Assenza et al., 2017; Tecchio et al., 2018), investigating adjunct non-invasive therapies such as c-tDCS is justified. This highlights even further the need for reliable and predictive application of c-tDCS, with investigation into c-tDCS inter-individual variability aimed at ultimately optimising the number of responders and minimising the number of non-responders. By investigating potential intrinsic factors that may serve as predictive tools (i.e. genetic markers), the administration of c-tDCS may be allocated to those who will benefit the most. This theoretical framework combined with previous reports that specific variations in genes that encode for the inhibitory GABA receptors were associated with drugresistance in Epilepsy (Hung et al., 2013), highlight the need for investigation into the role of genetic variants in inter-individual variability to c-tDCS.

This study therefore aimed to be the first of its kind to investigate the relationship between NMDA and GABA receptor gene variants and inter-individual variability to c-tDCS. In a large sample-size, this study also aimed to investigate the predictive value of the selected gene variants for response to c-tDCS. The significance of this study will be its novelty in investigating c-tDCS, and its large-scale nature which will facilitate meaningful subgroup statistical analysis, thus increasing the power and impact of the research findings. This may provide insight into future studies investigating the application of c-tDCS whereby reductions in cortical excitability and CSE is desired such as in excitotoxic neurological populations such as Epilepsy and those prone to seizures.

We hypothesised there would be an association and predictive capacity between the selected gene variants and c-tDCS response. We hypothesised normal expression of genes encoding for GABA receptors would be associated with reductions in CSE and categorisation as c-tDCS responders while variations in GABA receptor genes would be associated with increases in CSE and c-tDCS non-responders. We also hypothesised that normal expression of NMDA receptor genes would be associated with c-tDCS non-responders while variant expression would be associated with c-tDCS responder categorisation. Changes in cortico-cortical excitability measures of intracortical inhibition (ICI) and intracortical facilitation (ICF) were also investigated and whether the relationship between c-tDCS response and ICI and ICF was different between responders and nonresponders.

## Methods

### Subjects

Monash University human research ethics committee granted approval of this study. Sixty-one healthy male volunteers aged (mean±SD) 26.82±7.62 years who had already participated in a previous tDCS study in the same lab (Pellegrini et al., 2020a), provided written informed consent to attend two experimental sessions. Sample-size (with 80% power) was based on pilot data of fifteen subjects (Pellegrini et al., 2020b). An effect size of 0.45 (α=0.05) required sample-sizes between 17-26 (Portney and Watkins, 2009). Sample size was adjusted to allow for responders and nonresponders. Previous literature report the proportion of c-tDCS responders range from 40-55%, with average responder rates approximately 47% (Strube et al., 2016; Wiethoff et al., 2014). To ensure the numbers of c-tDCS responders were between 17-26, sample-size was adjusted to at least fifty-seven (i.e. 26/0.47=57). Female participants were not recruited to maximise subject homogeneity and control for effects of fluctuating estrogen and progesterone hormones (Inghilleri et al., 2004; Smith et al., 2002; Zoghi et al., 2015). Fifty-four subjects were right-handed and seven left-handed as determined by the Edinburgh handedness questionnaire (Oldfield, 1971) and no subject reported neurological or psychological conditions (Keel et al., 2001). All subjects refrained from consuming caffeine at least 12-hours prior to experimental testing (Chew et al., 2015; Fujiyama et al., 2017; Hermsen et al., 2016; Matamala et al., 2018; O’leary et al., 2015) to minimise confounding effects of caffeine (Cerqueira et al., 2006; Concerto et al., 2017).

### Study Design

A repeated-measures randomised cross-over design was utilised. Subjects attended two identical sessions in randomised order (c-tDCS or sham-tDCS). Sessions were conducted at similar times-of-day to reduce cortisol diurnal effects (Sale et al., 2008, 2007) and separated by at least one-week ensuring no carry-over effects (Nitsche et al., 2008).

### Electromyography Recording

Electromyography (EMG) was recorded for the dominant first dorsal interossei (FDI). Skin was abraded and cleaned to minimise skin impedance (Gilmore and Meyers, 1983). Pre-gelled self-adhesive bipolar Ag/AgCl disposable surface electrodes were used and placed over the FDI with 2cm inter-electrode distance and reference electrode over ulna styloid process (Kendell et al., 2010). EMG signals were filtered, amplified (10-500Hz x 1000) and sampled at 1000Hz and collected via a laboratory analogue-digital interface (LabChart™ & Powerlab, ADInstruments, Australia).

### Intra-Rater Reliability for assessment of corticospinal excitability

Single-assessor (MP) intra-rater reliability for the assessment of corticospinal excitability via TMS has been previously established (Pellegrini et al., 2018c). CSE, as measured by peak-to-peak amplitude of MEPs, were recorded in several TMS test intensities; 105%, 120%, 135%, 150%, 165% of resting motor threshold (RMT). Significant inter-class correlations ranging from 0.660-0.968 (p<0.05) were reported both within-sessions and between-sessions for TMS test intensities 120-165% of RMT (Pellegrini et al., 2018c).

### Transcranial Magnetic Stimulation Procedure

Single and paired-pulse stimuli were delivered to the dominant primary motor cortex (M1) by a 70mm figure-of-eight TMS coil (Magstim Limited Company, UK). Held over M1, the coil was oriented 45° to the midline and tangential to the scalp for posterior-anterior current flow (Rossini and Rossi, 1998) and repositioned to determine the cortical area eliciting the greatest FDI motor response. This hotspot was marked for consistent coil placement.

RMT was defined as the percentage of TMS device maximal stimulator output (MSO) required for MEP peak-to-peak amplitude greater than 50μV in 5/10 consecutive stimuli (Devanne et al., 2006). Test intensity was defined as the percentage of TMS MSO required to elicit an MEP peak-to-peak amplitude of ~1mV. The intensity was adjusted in 1-2% intervals until RMT and test intensity were determined (Rothwell et al., 1999). These were re-calculated post-tDCS intervention for each session.

### Outcome Measures

To assess CSE, twenty-five MEPs at the test intensity with 6sec inter-trial interval (ITI) were recorded. Cortico-cortical excitability were assessed via TMS paired-pulse paradigms. A conditioning pulse at 80% RMT followed by a pulse at the test intensity separated by 3msec inter-stimulus interval (ISI) for short-interval intracortical inhibition (SICI) and 10msec for ICF (Di Pino et al., 2014; Kujirai et al., 1993). Fifty paired-pulse MEPs (25 with 3msec ISI, 25 with 10msec ISI) were delivered with 6sec ITI. Mean values were calculated for SICI and ICF, then expressed as a percentage of single-pulse MEPs, and were considered as an index of M1 ICI and ICF (Di Pino et al., 2014; Kujirai et al., 1993).

### Transcranial Direct Current Stimulation

To maintain consistency with previous literature, tDCS was delivered at the common parameters of 1mA current intensity (Labruna et al., 2016; Strube et al., 2016) and 10-minute stimulus duration (Labruna et al., 2016; Strube et al., 2016; Wiethoff et al., 2014) with 30sec fade-in/fade-out periods (NeuroConn DC-stimulator, Germany). These parameters were also chosen to minimise the effect adjusting the stimulus parameters may have on responses, as adjusting stimulus parameters has been previously reported to influence responses to c-tDCS (Batsikadze et al., 2013). Two rectangular saline-soaked surface electrodes fixed to the scalp via velcro straps delivered the tDCS. The active electrode (5×7 cm, 0.0417mA/cm^2^) was placed over the contralateral supraorbital area and the return electrode (4×6cm, 0.0286mA/cm^2^) was placed over the dominant M1, focussing current under the anode and away from the cathode (Nitsche et al., 2007). For sham-tDCS, electrode placements were the same, with current increasing from 0-1mA for a 30sec fade-in period then reducing to 0 mA for the remaining 9.5-minutes. Participants were blinded to intervention, with blinding integrity assessed by asking participants on the nature of both interventions at the conclusion of both sessions.

While considered a safe intervention (Nitsche et al., 2008), tDCS tolerability and side-effects were assessed during and after tDCS was administered. Sensations of itching, tingling or discomfort were monitored throughout the application of tDCS via participants being asked to rate the presence of sensations on a scale of 1-10 at the beginning and middle of tDCS application. The presence of headache or other sensory complaints following tDCS were also assessed on a scale of 1-10.

### Genotyping Procedure

Oragene-DNA self-collection kits (DNAgenotek, Ontario, Canada) were used to obtain the saliva samples. Ten genetic single nucleotide polymorphisms (SNP) were chosen based on a previous similar study investigating the role of genetic polymorphisms in inter-individual variability to tDCS (Pellegrini et al., 2020a). The ten SNPs were selected as they are involved in synaptic transmission. SNPs for the BDNF gene (Antal et al., 2010; Cheeran et al., 2008; Di Lazzaro et al., 2015; Frazer et al., 2016; Puri et al., 2015) and glutamate NMDA receptor genes GRIN1 (Lee et al., 2016; Mori et al., 2011; Rossi et al., 2013) and GRIN2B (Mori et al., 2011; Narita et al., 2018) were selected based on their involvement in excitatory glutamatergic cortical pathways. GABA receptor genes GABRA1, GABRA2 and GABRA3 (Hung et al., 2013) were selected for their involvement in inhibitory GABAergic cortical pathways (table 1). Genotyping was performed by the Australian Genome Research Facility (AGRF, St Lucia, Australia) once all data collection of all participants were completed to avoid assessor bias. Subjects classification was either ‘normal expression’ (i.e. common homozygote) or ‘variant expression’ (i.e. heterozygous or homozygous for the substituted nucleotide) for each of the selected gene SNPs.

**Table 1.** Selected genes and SNPs. Summary of selected genes and SNPs with normal expression (common homozygous) and variant expression (heterozygous and rare homozygous) with accompanying sample-sizes.

| Gene | Protein/Receptor | SNP | Common |  | Variant |  |
| --- | --- | --- | --- | --- | --- | --- |
|  |  |  | Expression | N | Expression | N |
| BDNF | Brain-derived Neurotrophic Factor | rs6265 | CC | 31 | CT/TT | 30 |
| GRIN1 | Glutamate NMDA receptor subunit 1 | rs6293 | AA | 37 | AG/GG | 24 |
|  |  | rs4880213 | CC | 24 | CT/TT | 37 |
| GRIN2B | Glutamate NMDA receptor subunit 2 | rs1805247 | AA | 53 | AG/GG | 7 |
|  |  | rs7301328 | GG | 22 | GC/CC | 39 |
| GABRA1 | GABA receptor subunit $\alpha 1$ | rs6883877 | CC | 51 | CT/TT | 3 |
| GABRA2 | GABA receptor subunit $\alpha 2$ | rs279871 | TT | 23 | TC/CC | 31 |
|  |  | rs511310 | GG | 46 | GA/AA | 15 |
| GABRA3 | GABA receptor subunit $\alpha 3$ | rs1112122 | GG | 30 | GT/TT | 30 |
|  |  | rs4828696 | TT | 31 | TC/CC | 29 |

### Experimental Procedure

The sessions (c-tDCS, sham-tDCS) were conducted in randomised order. Subjects sat relaxed in an adjustable chair with their hand at rest. RMT and test intensities were determined and then baseline measures were collected. One of the two tDCS interventions were then delivered and immediately followed by CSE, via single-pulse MEP, outcome measure data collection at 0-minutes post-tDCS. RMT and test intensities were then re-established for SICI and ICF data collection at 10-minutes post-tDCS. Single-pulse MEPs were collected at 30-minutes post-tDCS using the baseline intensity while SICI and ICF were collected at 40-minutes post-tDCS using the adjusted intensity to investigate whether outcome measures had returned to baseline values (A. Bastani and Jaberzadeh, 2013; Andisheh Bastani and Jaberzadeh, 2013). Saliva samples were collected at the conclusion of the second session.

### Statistical Analyses

Statistical analyses were conducted using SPSS (Version 25.0, IL, USA).

#### Group level analysis

Each outcome measure (CSE, SICI, ICF) was assessed for each intervention (c-tDCS, sham-tDCS) and time-point (baseline, 0-10 minutes, 30-40 minutes). Shapiro-Wilk test assessed data normality for each outcome measure. Non-parametric statistical tests were used if data violated normality.

Comparison of baseline values for each session assessed the effect of intervention on within-subject reliability to ensure no carry-over effects between sessions (Portney and Watkins, 2009). Non-parametric Friedman’s two-way analysis of variance (ANOVA) was used to investigate the overall effect of time and intervention on CSE, SICI and ICF data which violated normality. Non-parametric Wilcoxon matched-paired test was then used post-hoc where appropriate to investigate whether outcome measures at post-tDCS time-points were significantly different from baseline values following both c-tDCS and sham-tDCS. 5% significance level was used for all statistical tests (α=0.05).

#### Subgroup level analysis

Two subgrouping techniques were used to categorise subjects as defined and described in detail previously (Pellegrini et al., 2018b). Firstly, subjects were categorised based on their post-tDCS responses exceeding a pre-determined thresholds or not. Normalised to baseline, MEP amplitudes at 0-minutes post-tDCS were compared to the pre-determined threshold of the standard deviation (SD) of sham-tDCS baseline values (Ammann et al., 2017). This accounted for natural response variability in the current sample size as well as the inherent variability in measuring CSE via TMS-evoked MEPs. In this current study, using a grand average of all post-tDCS time-points as previously conducted (Ammann et al., 2017; Chew et al., 2015; López-Alonso et al., 2014; Puri et al., 2016, 2015; Tremblay et al., 2016; Wiethoff et al., 2014) would not have provided an inaccurate representation of how each subject immediately responded to c-tDCS because MEP data collection at 30-minutes post-tDCS has been reported to return to baseline levels (A. Bastani and Jaberzadeh, 2013; Andisheh Bastani and Jaberzadeh, 2013). Sham-tDCS baseline SD calculated at 0.324, therefore classified subjects as ‘responders’ (post c-tDCS nMEP<0.676) or ‘non-responders’ (post c-tDCS nMEP>0.676).

The second subgrouping technique was the SPSS TwoStep cluster analysis. Based on normalised data, analysis of trends or patterns in the data highlights distinct subgroups of individuals (Hair et al., 1998; SPSS, 2001). Cluster predictors were normalised MEP amplitude at 0-minutes post c-tDCS.

Non-parametric Mann-Whitney U test was conducted on MEP data to investigate significant differences between the two distinct subgroups identified (i.e. repsonders/non-responders and cluster1/cluster 2).

Subgroups were then analysed separately to investigate the effect of time and intervention on SICI and ICF. Non-parametric Friedman and Wilcoxon matched-paired tests were used to investigate whether there were significant changes in SICI and ICF at post-tDCS time-points compared to baseline following both c-tDCS and sham-tDCS for each subgroup. Non-parametric Mann-Whitney U tests were then conducted to determine significant differences between subgroups (i.e. responders versus non-responders or cluster 1 versus cluster 2) at each time-point.

#### Genotype analysis

Genotype results were analysed against both subgroup categorisation techniques for investigation of associations between gene expression and c-tDCS response categorisation. This was to test whether gene expression had predictive capacity for c-tDCS response category. Chi-squared tests analysed each gene individually against subgroup category and cluster membership tested for associations/dependency between gene expression and c-tDCS response. Univariate binary logistical regression analysis then quantified the association (i.e. odds ratio) between each gene variant independently and subgroup category to assess predictive value of each gene variant in isolation for c-tDCS response. Dependent variable was the subgroup category and each gene was the independent variable. Multivariate binary logistical regression analysis was then conducted as above to investigate associations and predictive value of each gene variant for c-tDCS response while controlling for all other gene variants.

## Results

All sixty-one participants attempted each session with mean (±SD) 13.58±15.91 days between each session. Mean RMT and test intensity were 37% (36.55±7.59%) and 46% (45.58±9.88%) of the TMS MSO.

### Tests for Normality of data

Shapiro-Wilk tests for normality revealed single-pulse MEP, SICI and ICF data violated normality (p<0.05).

### Carry-Over effects

Friedman’s test revealed no significant differences in baseline values between c-tDCS and sham-tDCS for single-pulse MEP (p=0.370), SICI (p=0.249) or ICF (p=0.701) indicating no carry-over effects between sessions.

### Group level analysis

#### Corticospinal Excitability

Non-parametric Friedman’s test and post-hoc Wilcoxon matched-paired test revealed no significant effect of time for both c-tDCS and sham-tDCS (p>0.05) but a significant effect of intervention with MEP amplitude significantly reduced at 0-minutes following c-tDCS compared to the same timepoint following sham-tDCS (p<0.01). Differences in MEP between interventions at 30-minutes were not significantly different (p>0.05) (figure 1a-b and table 2).

**Figure 1.**
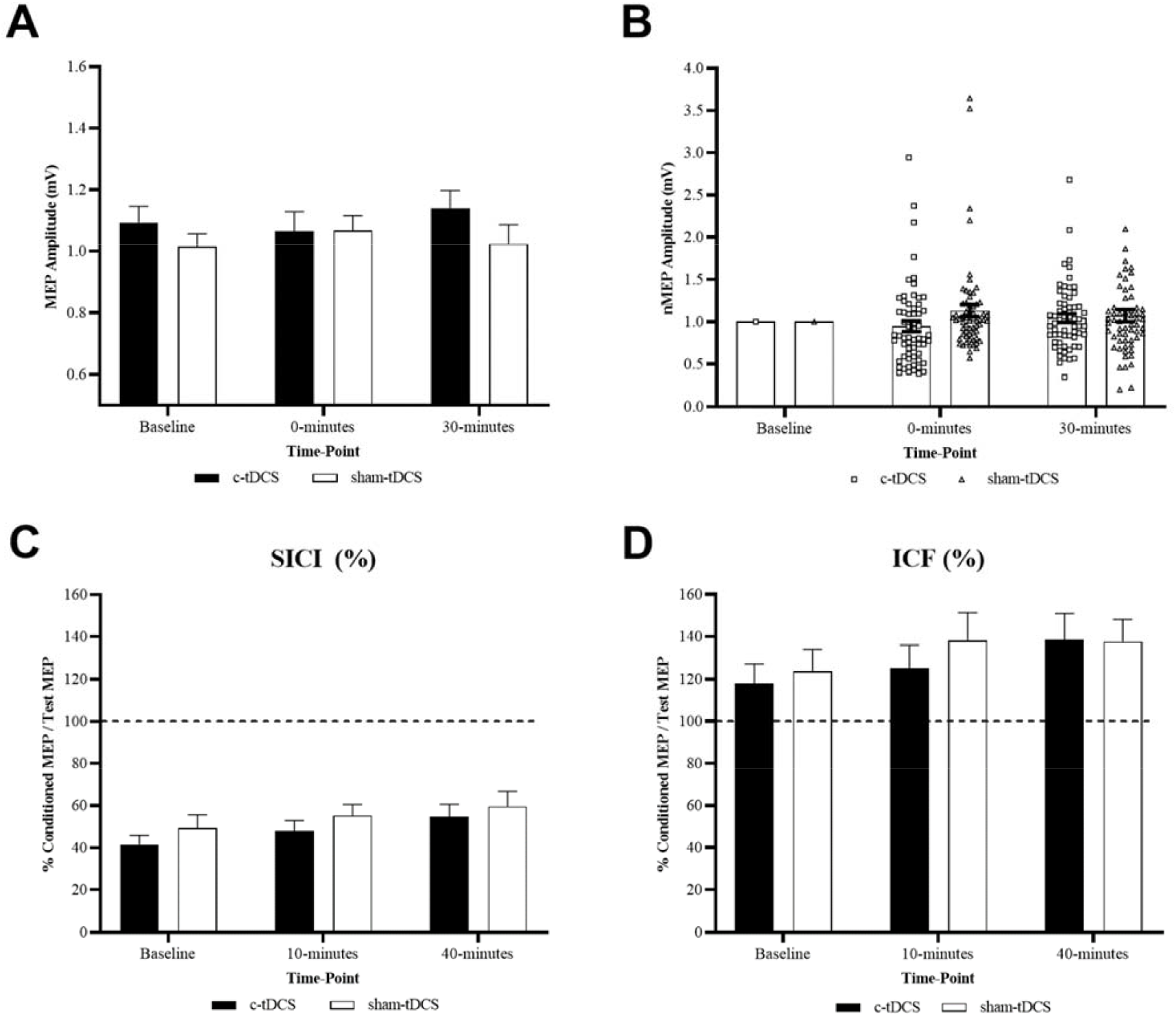
Group level c-tDCS versus sham-tDCS for each time-point. (a) Mean single-pulse MEP amplitudes. (b) Changes in single-pulse MEP amplitude normalised to baseline with individual data points. (c) ICI as measured by SICI values. (d) ICF. Error bars denote one SEM.

**Table 2.** Group level corticospinal excitability and cortico-cortical excitability. Mean(±SD) MEP peak-to-peak amplitude (mV), SICI (%) and ICF (%) data for each intervention and time-point. Post-tDCS time-point 1 is 0-minutes for MEP and 10-minutes for SICI and ICF. Post-tDCS time-point 2 is 30-minutes for MEP and 40-minutes for SICI and ICF. ^*^denote significant difference in MEP amplitude between interventions at a particular time-point (p<0.01).

| **Intervention** | **Outcome Measure** | **Baseline** | **Post-tDCS time-point 1** | **Post-tDCS time-point 2** |
| --- | --- | --- | --- | --- |
| **c-tDCS** | **MEP (mV)** | 1.102±0.340 | 0.996±0.490* | 1.120±0.433 |
| | **SICI (%)** | 41.60±33.53% | 47.89±39.41% | 54.76±45.54% |
| | **ICF (%)** | 117.88±73.27% | 125.19±85.18% | 138.58±97.21% |
| **sham-tDCS** | **MEP (mV)** | 1.015±0.324 | 1.066±0.377* | 1.024±0.487 |
| | **SICI (%)** | 49.28±49.78% | 55.22±41.02% | 59.67±55.50% |
| | **ICF (%)** | 123.71±81.01% | 138.13±103.22% | 137.66±81.51% |

#### Cortico-cortical Excitability

Non-parametric Friedman’s test and post-hoc Wilcoxon matched-paired test revealed no significant effect of time or intervention for SICI and ICF for both active and sham c-tDCS (p>0.05) (figure 1c-d and table 2).

### Individual level analysis

#### Subgrouping based on a pre-determined threshold

Subgrouping based on normalised responses at 0-minutes post c-tDCS with a 32.4% threshold revealed two different subgroups. Following c-tDCS, 30% (n=18) of subjects responded with reductions in MEP that exceeded the predetermined threshold of 0.324 and were subsequently categorised as ‘responders’ (nMEP<0.676 at 0-minutes post c-tDCS) while 70% (n=43) responded with no reductions or even increases in MEP and were categorised as ‘non-responders’ (nMEP>0.676 at 0-minutes post c-tDCS) (figures 2c).

**Figure 2.**
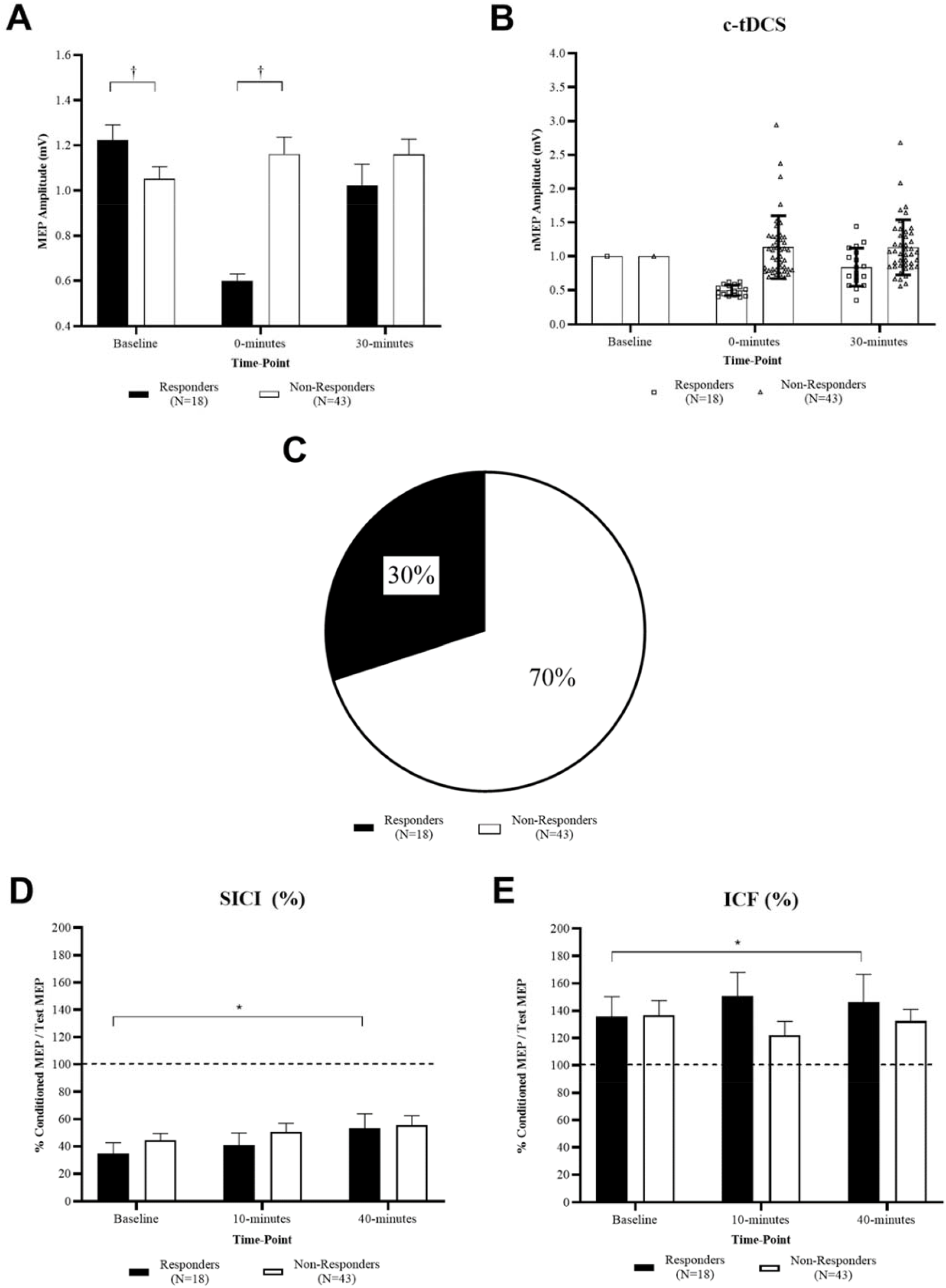
Subgrouping based on pre-determined threshold. Responders versus Non-Responders before and after c-tDCS at each time-point. (a) Single-pulse MEP amplitudes. (b) Single-pulse normalised MEP amplitude with individual data-points. (c) Percentage breakdown of responders and non-responders. (d) ICI as measured by SICI. (e) ICF. Error bars denote one SEM. ^†^denote significant differences between responder and non-responder subgroups at a particular time-point (p<0.05). ^*^denote significant difference between time-points (p<0.05).

##### Corticospinal Excitability

Mann-Whitney U tests revealed MEP amplitudes were significantly different between responder and non-responder subgroups for c-tDCS at baseline and 0-minutes post-tDCS (p<0.05) but not at 30-minutes post-tDCS (p>0.05). No differences between responders and non-responders for sham-tDCS were reported (p>0.05). Additionally, for responders, MEP values did not differ between c-tDCS and sham-tDCS interventions at baseline or 30-minutes post-tDCS (p>0.05) but were significantly different at 0-minutes post-tDCS (p<0.05). For non-responders, MEP were not significantly different between interventions at baseline or 0-minutes post-tDCS (p>0.05) but were at 30-minutes post-tDCS (p<0.05) (figure 2a-c and table 3).

**Table 3.**
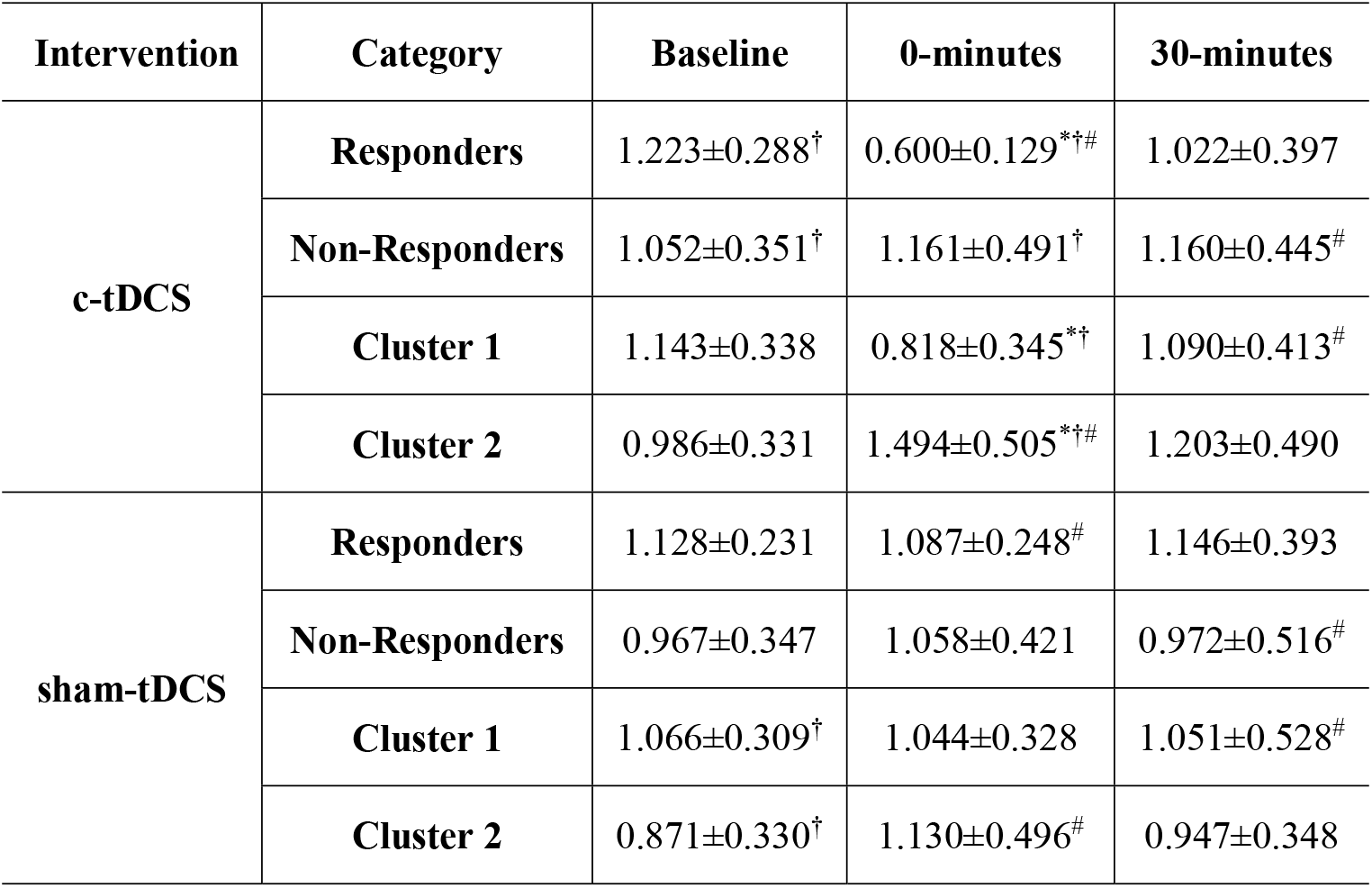
Corticospinal excitability for responders/non-responders and cluster 1/cluster 2 subgroups. Mean(±SD) MEP peak-to-peak amplitude (mV) for both interventions and time-point. ^*^denote significant difference within a subgroup MEP amplitude at a time-point compared to baseline (p<0.05). ^†^denote significant difference in MEP amplitude at a particular time-point between responder and non-responder subgroups as well as between cluster 1 and cluster 2 subgroups (p<0.05). ^#^denote significant difference between tDCS interventions for a particular subgroup (p<0.05).

##### Cortico-cortical Excitability

For SICI, in the responder subgroup non-parametric Friedman’s test revealed a significant effect of time for c-tDCS, with post-hoc Wilcoxon matched-paired tests revealing significant increases in SICI values, indicating significant reductions in ICI at 30-minutes following c-tDCS compared to baseline (p<0.05). No significant differences were reported between interventions for the responder subgroup. For sham-tDCS, no significant effect of time or intervention were reported for both responder and non-responder subgroups (p>0.05). Mann-Whitney U test revealed no significant differences between subgroups at all time-points for both c-tDCS and sham-tDCS (p>0.05) (figure 2d and table 4).

**Table 4.** Cortico-cortical excitability for responder and non-responder subgroups. Mean(±SD) SICI and ICF data for each intervention and time-point. ^*^denote significant difference within a subgroup at a time-point compared to baseline (p<0.05). ^†^denote significant difference at a particular time-point between responder and non-responder subgroups (p<0.05).

| Outcome Measure | Subgroup | SICI (%) |  |  | ICF (%) |  |  |
| --- | --- | --- | --- | --- | --- | --- | --- |
|  |  | Baseline | 10-minutes | 40-minutes | Baseline | 10-minutes | 40-minutes |
| c-tDCS | Responder | 34.93±32.88% | 41.00±37.44% | 53.21±44.98%* | 74.09±26.01%† | 90.64±31.85% | 111.84±33.47%* |
| | Non-Responder | 44.39±33.79% | 50.78±40.28% | 55.40±46.28% | 136.22±78.92%† | 139.65±96.06% | 149.77±112.30% |
| sham-tDCS | Responder | 41.01±23.04% | 50.42±34.62% | 52.24±30.79% | 91.89±40.43%† | 116.69±68.59% | 111.55±40.25% |
| | Non-Responder | 52.74±57.30% | 57.23±43.64% | 62.79±63.11% | 137.03±89.97%† | 147.10±114.18% | 148.59±91.76% |

For ICF, in the responder subgroup a significant effect of time for c-tDCS, with post-hoc Wilcoxon matched-paired tests revealed significant increases in ICF at 40-minutes following c-tDCS compared to baseline (p<0.05) while no significant differences were reported between interventions for the responder subgroup. For sham-tDCS, no significant effect of time or intervention were reported for both subgroups (p>0.05). Mann-Whitney U test revealed no significant differences between responder and non-responder subgroups at each time-point and intervention (p>0.05) (figure 2e and table 4).

#### Two-Step Cluster Analysis

Subgrouping based on SPSS statistical cluster analysis revealed two distinct subgroups. Cluster 1 comprised 74% (n=45) of subjects that responded with an overall average reduction in MEP. Cluster 2 comprised 26% (n=16) of subjects that responded with either no reduction or overall increase in MEP (figure 3c).

**Figure 3.**
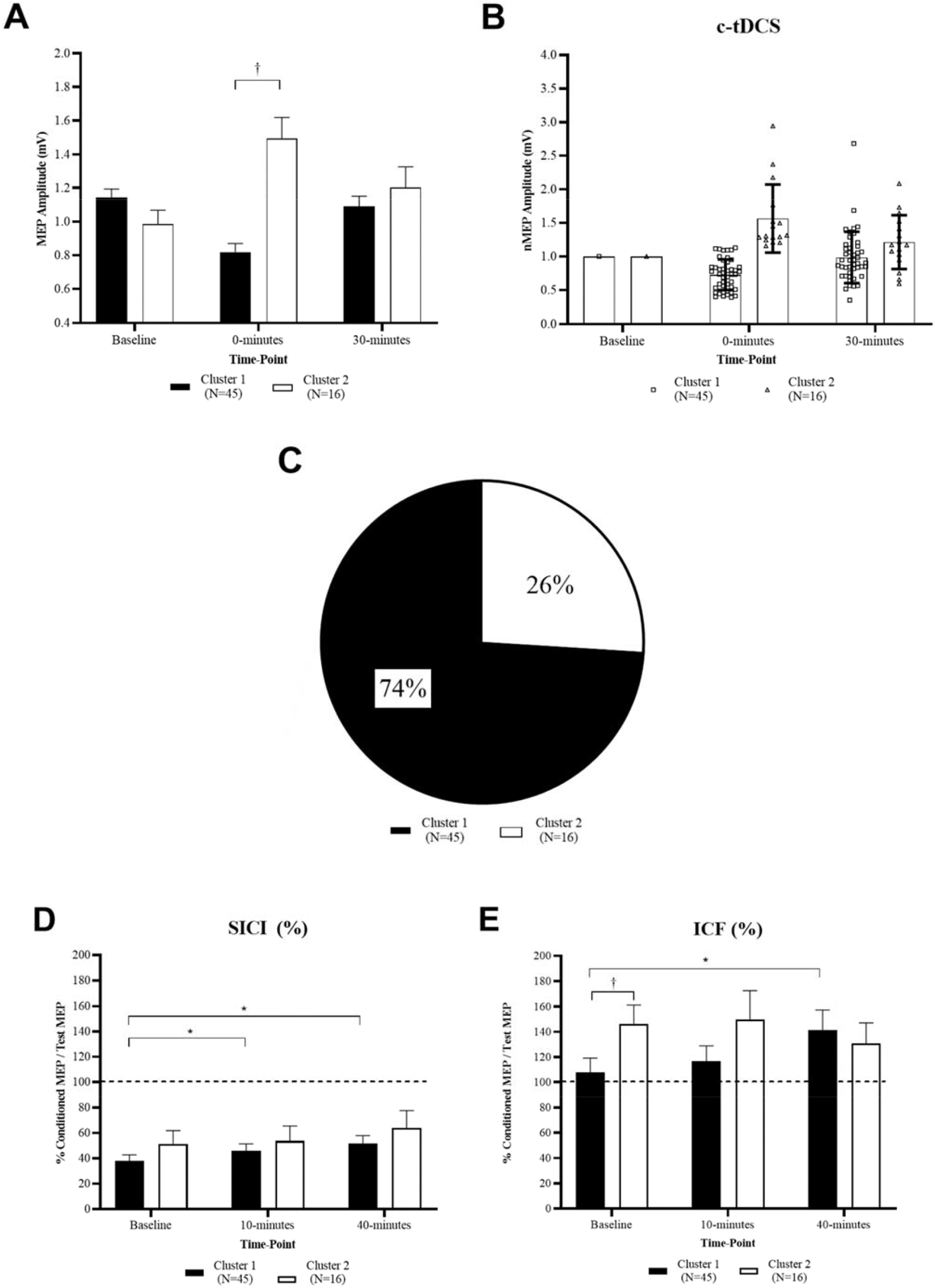
Subgrouping based on cluster analysis. Cluster 1 (responders) versus cluster 2 (non-responders) before and after c-tDCS at each time-point. (a) Single-pulse MEP amplitudes. (b) Single-pulse normalised MEP amplitude with individual data-points. (c) Percentage breakdown of cluster 1 (responders) and cluster 2 (non-responders). (d) ICI as measured by SICI. (e) ICF. Error bars denote one SEM. ^†^denote significant differences between cluster 1 and cluster 2 subgroups at a particular time-point (p<0.05). ^*^denote significant difference between time-points (p<0.05).

##### Corticospinal Excitability

For c-tDCS, Mann-Whitney U tests reported no significant differences in MEP amplitude between cluster 1 and cluster 2 subgroups at baseline and 30-minutes post-tDCS (p>0.05), but revealed MEP amplitude was significantly greater in cluster 2 at 0-minutes following c-tDCS compared to cluster 1 (p<0.05). Significant differences were reported between subgroups for sham-tDCS at baseline (p<0.05), but not for 0-minutes and 30-minutes post-tDCS (p>0.05). For cluster 1, no significant differences in MEP amplitude were reported between c-tDCS and sham-tDCS at baseline and 30-minutes (p>0.05) but were reported at 0-minutes post-tDCS (p<0.05). For cluster 2, significant differences were reported between c-tDCS and sham-tDCS at 30-minutes (p<0.05) but not at baseline or 0-minutes post-tDCS (p>0.05) (figure 3a-c and table 3).

##### Cortico-cortical Excitability

For SICI, in the cluster 1 subgroup, Friedman’s test and post-hoc Wilcoxon matched-paired tests revealed significant increases in SICI values at 10-minutes and 40-minutes following both c-tDCS and sham-tDCS compared to baseline (p<0.05), indicating significant reductions in ICI. For the cluster 2 subgroup non-parametric Friedman’s test revealed no significant effect of time or intervention following both c-tDCS and sham-tDCS (p>0.05). For clusters 1 and 2, no significant differences were reported between interventions for both c-tDCS and sham-tDCS (p>0.05). Mann-Whitney U tests revealed no significant differences in SICI between cluster subgroups for both c-tDCS and sham-tDCS at all time-points (p>0.05) (figure 3d and table 5).

**Table 5.**
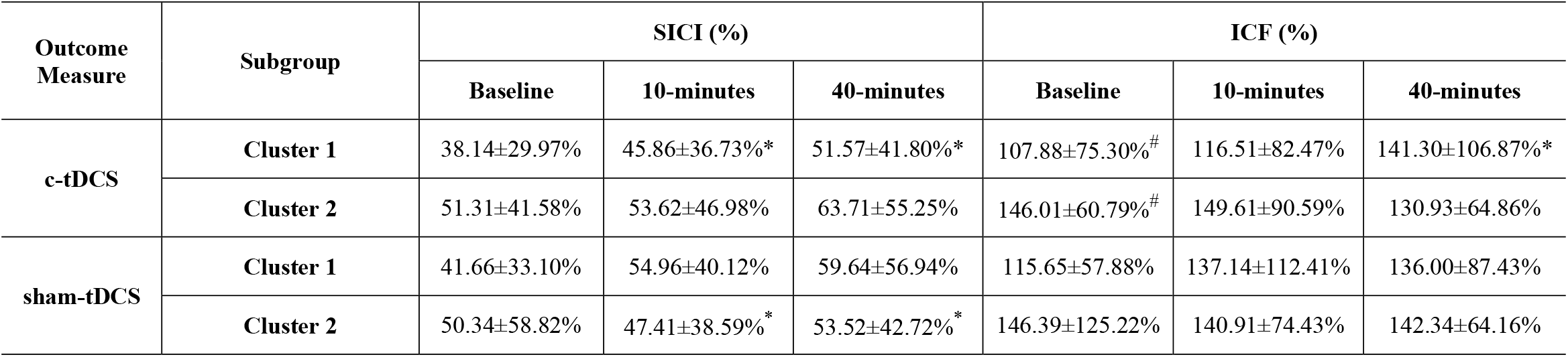
Cortico-cortical excitability for cluster 1 and cluster 2 subgroups. Mean(±SD) SICI and ICF data for each intervention and time-point. ^*^denote significant difference within a subgroup at a time-point compared to baseline (p<0.05). ^#^denote significant difference between tDCS interventions for a particular subgroup at a particular time-point (p<0.05).

For ICF, in the cluster 1 subgroup, Friedman’s test and post-hoc Wilcoxon matched-paired tests reported no significant changes in ICF at 10-minutes following c-tDCS compared to baseline but revealed significant increases at 40-minutes following c-tDCS (p<0.05). For the cluster 2 subgroup non-parametric Friedman’s test revealed no significant effect of time or intervention following both c-tDCS (p>0.05). No changes were reported in cluster 1 and 2 subgroups at all sham-tDCS timepoints (p>0.05). No differences were reported between interventions at each time-point for both cluster 1 and 2 subgroups (p>0.05). Mann-Whitney U tests revealed ICF values were significantly greater at c-tDCS baseline in the cluster 2 subgroup compared to cluster 1 (p<0.05). No other significant differences were reported between cluster subgroups for c-tDCS, or at all sham-tDCS time-points (p>0.05) (figure 3e and table 5).

### Genotype analysis

#### Association/dependency between gene expression and inter-individual variability

Genotyping of this cohort of subjects (n=61) revealed three of the gene SNPs (i.e. GRIN2B rs1805247, GABRA1 rs6883877 and GABRA2 rs511310) had disproportionate sample-sizes in each dichotomous subgroup (i.e. ‘normal expression’ and ‘variant expression’) to carry out comparative analysis and were therefore excluded from statistical analysis. Refer to table 1 for subgroup sample sizes.

Chi-squared analysis on the remaining seven gene SNPs revealed no significant associations between subgroup category based on pre-determined threshold or cluster analysis and genotype subgroup indicating dependency between expression of the selected genes and responder/non-responder or cluster 1/cluster 2 subgroups (table 6).

**Table 6.**
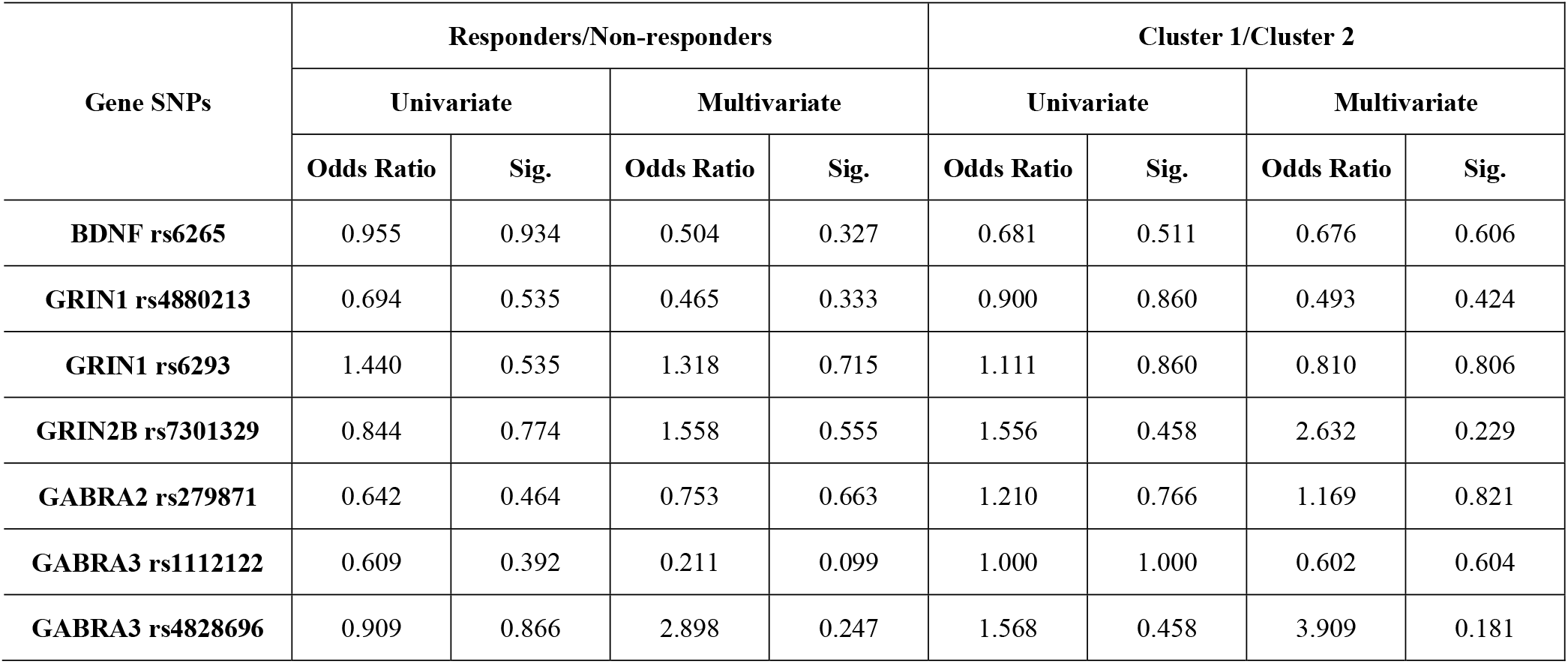
Odds ratio and logistical regression analysis results comparing each of the selected gene SNPs with both responder/non-responder and cluster 1/cluster 2 subgroups.

| Gene SNPs | Responders/Non-responders |  |  |  | Cluster 1/Cluster 2 |  |  |  |
| --- | --- | --- | --- | --- | --- | --- | --- | --- |
|  | Univariate |  | Multivariate |  | Univariate |  | Multivariate |  |
|  | Odds Ratio | Sig. | Odds Ratio | Sig. | Odds Ratio | Sig. | Odds Ratio | Sig. |
| **BDNF rs6265** | 0.955 | 0.934 | 0.504 | 0.327 | 0.681 | 0.511 | 0.676 | 0.606 |
| **GRIN1 rs4880213** | 0.694 | 0.535 | 0.465 | 0.333 | 0.900 | 0.860 | 0.493 | 0.424 |
| **GRIN1 rs6293** | 1.440 | 0.535 | 1.318 | 0.715 | 1.111 | 0.860 | 0.810 | 0.806 |
| **GRIN2B rs7301329** | 0.844 | 0.774 | 1.558 | 0.555 | 1.556 | 0.458 | 2.632 | 0.229 |
| **GABRA2 rs279871** | 0.642 | 0.464 | 0.753 | 0.663 | 1.210 | 0.766 | 1.169 | 0.821 |
| **GABRA3 rs1112122** | 0.609 | 0.392 | 0.211 | 0.099 | 1.000 | 1.000 | 0.602 | 0.604 |
| **GABRA3 rs4828696** | 0.909 | 0.866 | 2.898 | 0.247 | 1.568 | 0.458 | 3.909 | 0.181 |

Univariate logistic regression analysis and odds ratio (OR) calculation revealed no significant predictive value for each of the selected gene SNPs when analysed in isolation for either subgroup categorisation based on pre-determined threshold (i.e. responder/non-responders) or cluster membership (p>0.05). Refer to table 6 for OR values for each of the selected SNPs in isolation.

When all of the selected SNPs were controlled and accounted for, multivariate logistic regression analysis with OR calculation again revealed no significant predictive value of each of the selected SNPs for response subgroup categorisation (p>0.05). It is worth noting however that subjects with GABRA3 gene SNP rs1112122 ‘normal expression’ were five times (OR=4.746) more likely to be categorised as a c-tDCS responder based on pre-determined threshold compared to subjects with ‘normal expression’, with ‘variant expression’ subjects less likely (OR=0.211) to be categorised as a c-tDCS responders. These results were approaching significance (p=0.099). Therefore when each of the selected gene SNPs were controlled and accounted for, no significant associations suggest no predictive value for subgroup response category (table 6).

## Discussion

This study investigated response variability to c-tDCS and the role genetic variations played in the observed inter-individual variability. We hypothesised that variations in genes encoding for excitatory NMDA and inhibitory GABA receptors would influence observed inter-individual variability to-tDCS. Two distinct subgroups of individuals were determined via two different subgrouping techniques. Both techniques yielded a group of individuals who responded as expected with reductions in CSE (i.e. responders) and another group who responded with no reduction or even increases in CSE (i.e. non-responders). Changes in both ICI and ICF were reported in both responder and non-responder subgroups. No significant relationships however were identified between c-tDCS response and the expression of both NMDA and GABA receptor genes.

### Group and subgroup level analysis

A sample size of sixty-one, to our knowledge is the largest in the tDCS literature that investigates c-tDCS inter-individual variability. Consistent with previous large-scale c-tDCS studies (Strube et al., 2016; Wiethoff et al., 2014), no overall effects were reported in the entire cohort but distinct subgroups were reported. Building upon the results from previous studies investigating interindividual variability to c-tDCS, the subgrouping criteria implemented in this current study aimed to take into account the natural variation that may occur when assessing CSE via TMS-evoked MEPs by setting the response threshold as the SD of the baseline sham-tDCS condition (Ammann et al., 2017). The resultant c-tDCS responder rate of 30% is considerably lower than the 53% (Strube et al., 2016) and 41% (Wiethoff et al., 2014) as previously reported. The discrepancy between our current results and previous studies may be a reflection of the more strict and controlled subgrouping threshold specific to our current cohort of individuals. The individual data points normalised to baseline in figure 2b provide a visual representation of the spread of responses between the sixty-one participants with a considerable number of individuals responding to c-tDCS with reductions in CSE that did not exceed our predetermined threshold of 32.4%. Having a more generalised subgrouping criteria of whether an individual increases or decreased CSE, that being whether their post c-tDCS normalised responses were greater than or less than 1mV (Strube et al., 2016; Wiethoff et al., 2014), would have yielded a higher percentage of responders in the current study, but may not have been a true reflection of responses to c-tDCS when inherent variability in the assessment tool is not considered.

Subgrouping individuals via statistical cluster analyses did not support the findings when subgrouping individuals based on a pre-determined threshold. In comparison to the 30% of individuals that responded to c-tDCS with reductions in CSE that exceeded the pre-determined threshold, 74% of the individuals were categorised into a cluster group that reported overall reductions in CSE (figure 3b). This percentage is considerably greater than that reported in previous studies of 51% (Strube et al., 2016) and 47% (Wiethoff et al., 2014). Reasons for this discrepancy with not only previous literature, but also with the other utilised subgrouping technique, may again lie in the spread of the normalised data between all sixty-one individuals. Working on a similar concept, statistical cluster analyses scan the data for trends, patterns and thresholds that may divide the data into distinct groups or clusters (Hair et al., 1998; SPSS, 2001). A large proportion of the participating individuals responded with small magnitude changes in CSE, as displayed by the large grouping of responses around 1mV at 0-minutes following c-tDCS (figure 3b). Given cluster analyses divide data into clusters with no priori information regarding the expected responses, in this current dataset the cluster analysis prioritised those individuals that had large responses in the opposite direction to expected (i.e. large increases in CSE) and were therefore considered a distinct cluster. While the two selected subgrouping techniques did not report subgroups of similar sample-sizes with similar magnitude changes in CSE, the wide-ranging magnitudes of responses add further weight to the presence of inter-individual variability to c-tDCS and highlight a need for investigating factors which may contribute.

One such investigation involved reporting whether or not the observed changes in CSE, and the differences between the dichotomous subgroups of responders and non-responders could in-part be explained by measures of cortico-cortical excitability. To our knowledge, this was the first study of its kind to investigate inter-individual variability to c-tDCS and also compare changes in cortico-cortical excitability measures ICI and ICF in responder and non-responder subgroups. No overall changes were reported in both ICI, as measured by SICI, and ICF following c-tDCS (figure 1c-d and table 2), but when subgrouped into responders and non-responders, significant changes were reported in both SICI and ICF at 40-minutes in the responder subgroup following c-tDCS with a trend toward facilitation, that being increased ICF and reduced ICI as indicated by increases in SICI values (figure 2c-d, figure 3c-d and tables 4-5). These significant changes at 40-minutes following c-tDCS as the trend toward facilitation in the responder subgroup may provide an explanation for the increases in CSE back towards baseline at the second post-tDCS time-point at 30-minutes (figure 2a-b and table 3). Despite these reported delayed changes in cortico-cortical excitability in the responder subgroup, no significant changes or differences in SICI or ICF values were observed between responder and non-responder subgroups at 0-minutes following c-tDCS that may explain the differences in CSE between the two subgroups at the same time-point. These results appear consistent with previously published studies investigating changes in cortico-cortical excitability following c-tDCS. Recent studies have reported no changes in SICI (Di Lazzaro et al., 2012; Sasaki et al., 2016) and ICF (Di Lazzaro et al., 2012; Vaseghi et al., 2016) following standard protocols of c-tDCS.

The results of this current study, may therefore suggest that inhibitory circuits, other than GABA-A receptor specific GABAergic circuits, which is measured via SICI, may explain the discrepancies in CSE between responders and non-responders following c-tDCS. This provides future opportunities to investigate other cortico-cortical excitability mechanisms to explain discrepancies in CSE between responders and non-responders immediately following c-tDCS. These may include investigating changes in long-interval intracortical inhibition (LICI), a measure of GABAergic inhibitory circuits involving GABA-B receptors (McDonnell et al., 2006) and short-interval afferent inhibition (SAI), a measurement of cholinergic circuitries that has been previously associated with changes in CSE following c-tDCS (Sasaki et al., 2016). Future studies investigating these different inhibitory circuitries may generate deeper insights into explanations behind inter-individual variability to c-tDCS.

### Genetic analysis

In conjunction with a previously published study (Pellegrini et al., 2020a), this study is the first and largest of its kind to investigate inter-individual variability to c-tDCS and the involvement of specific variants in genes that encode for key regulators along excitatory and inhibitory cortical pathways. The novel findings that when all genetic variants were accounted for, a trending towards significance association exists between a GABRA3 gene variant and c-tDCS non-responder categorisation suggests a potential relationship between GABRA3 gene variants and reduced capacity for ICI and c-tDCS non-responder categorisation. A potential relationship that warrants further large-scale investigation.

With no previous studies in the tDCS literature investigating variants in genes that encode for GABA receptors, looking to Epilepsy and seizure literature for the influence of genetic variations may offer insight into role genetic variations play in cortico-cortical excitability and CSE changes following c-tDCS. This is particularly important for potential future application of c-tDCS, known to historically induce inhibition, on excitotoxic populations such as Epilepsy and those prone to seizures. Drug-resistance is considered common in Epileptic populations (Abou El Ella et al., 2018; Assenza et al., 2017; Hung et al., 2013; Naimo et al., 2019; Tecchio et al., 2018). Links between drug-resistance and particular gene variants (Baghel et al., 2016; Hung et al., 2013) offer insight into the role gene variants play in the activity of inhibitory cortical pathways.

Specific to the selected gene variants in this current study, variants in the GABRA3 gene, responsible for encoding GABA-A receptors, have been associated with antiepileptic drug-resistance in Epilepsy populations (Hung et al., 2013). Gene variants associated with resistance to drugs that aim to reduce cortical excitation suggest the resultant GABA receptor activity and overall GABAergic inhibitory regulation of cortical excitability may be impacted. In this current study, an association approaching significance, that-being those with GABRA3 gene variants less likely to respond to c-tDCS as expected with reductions in CSE, suggests a potential reduced capacity for GABAergic inhibitory regulation in individuals categorised as c-tDCS non-responders. This potential relationship is consistent with recently published results suggesting a link between variants in genes that encode for GABA receptors and an increased likelihood of responding to excitatory a-tDCS with increases in CSE (Pellegrini et al., 2020a).

These novel preliminary suggestions of potential predictive value of genetic variants to tDCS inter-individual variability should become a focus of future large-scale research. Larger sample-sizes should be recruited to facilitate analysis of the interactions between a number of gene variants and subsequent responses to tDCS. This is with the ultimate aim to identify a predictive genetic marker or series of genetic markers to detect likely tDCS responders, that may be developed into a point-of-care screening tool for suitability to tDCS in the clinical setting.

### Limitations

In an attempt to minimise potential sources of inter-individual variability, the cohort of participants recruited was strictly specified as health young males. These attempts however do limit the capacity to extrapolate our findings to the wider community that includes females, as well as older and pathological populations. In additional attempts to report findings that were relatable to similar previously published studies, conventional electrode montages were utilised with common stimulus parameters. It is therefore unclear whether our current results can be extrapolated to other forms of tDCS such as high-definition tDCS, or whether similar responder/non-responder rates would apply if tDCS was administered with different stimulus parameters. Lastly, the outcome measures for this study focussed on cortico-cortical and corticospinal measures and therefore the current results do not offer insight into the effect of c-tDCS on motor behavioural tasks, whether there is a difference in motor performance between responders and non-responders and whether any differences are genetically mediated. These listed limitations offer suggestions for future research into interindividual variability to c-tDCS.

### Future Directions

No relationships were reported between inter-individual variability to c-tDCS and genetic variations in the selected SNPs. While an association between the GABRA3 gene SNP rs1112122 was approaching significance (p=0.099), that suggested those individuals with a variation in this GABA receptor gene may be more likely to be c-tDCS non-responders, a non-significant association meant at this stage there was no relationship between specific genetic variations and response to c-tDCS. This highlights the opportunity for future research to investigate this potential relationship further to determine whether the c-tDCS response categorisation is genetically mediated. To achieve this, continuing the trend from this current study and recently published study (Pellegrini et al., 2020a), future studies should aim to recruit even larger sample sizes such that higher powered analysis can be conducted on larger genetic subgroups, and such that investigation into the interaction between a number of selected gene variants can be conducted. For example, while acknowledging an even larger sample size bears a large administrative burden, larger sample sizes will allow individuals to be subgrouped based on the presence or absence of multiple GABA receptor gene variants at once, or the presence of GABA receptor gene variants paired with the absence of excitatory NMDA receptor gene variants. This may provide insight into combinations of inherent genetic expressions that increase the likelihood of responding to c-tDCS as expected that may form the basis of a genetic screening and predictive tool for suitability to c-tDCS as an intervention in the clinical setting.

Furthermore, future studies investigating the role genetic expression has on c-tDCS responses and inter-individual variability should aim to investigate the role of gene variants in other key regulators of cortical plasticity as well as on differing cohort demographics. As discussed above, with changes in SICI and ICF offering no insight into the observed differences in CSE between responders and non-responders following c-tDCS, gene variants that encode for GABAergic and inhibitory cortical circuits other than GABA-A specific circuits should be investigated. This may provide additional insight and contribute to greater knowledge of the genetic component of cortical plasticity following c-tDCS and the differences between individuals who respond as expected and those who do not. Addressing the above mentioned limitations by extending these future studies of other genetic variants into other populations such as females, older adults and patient populations, particularly those prone to increased cortical excitability or excitotoxicity, should also form the basis of future research.

On a more general note, future studies on inter-individual variability to tDCS should continue on from the growing trend that not all individuals that participate in tDCS are the same. When investigating specific intrinsic factors that may contribute to inter-individual variability it is crucial that all other potential confounding factors such as age-range, medication-use, caffeine consumption and attentional focus are kept as consistent as possible between individuals such that one specific factor may be isolated to investigate its involvement in inter-individual variability. Future studies should therefore utilise the recently developed checklist (Pellegrini et al., 2020c) to ensure thorough methodological measures are taken in order to ensure inter-individual variability is minimised.

## Conclusions & Implications

The significance of this current study is it provides additional insight into the issue of interindividual variability to c-tDCS. It highlights that not all individuals will respond to c-tDCS in the same manner and therefore may not undergo cortical plasticity to the same extent when exposed to the same stimulus. While trending towards significance for one specific gene variant, the disparity in response between individuals did not appear to be genetically mediated. Implications of this variability in response for future studies are important when considering the clinical setting. For tDCS to be administered in the clinical setting in a large-scale manner, it must be a reliable, predictable and re-producible intervention. Studies such as this current study that systematically investigate one potential intrinsic factor and its role in resultant inter-individual variability are important steps forward for the development of a screening tool to determine those appropriate for c-tDCS interventions and who may benefit the most. Identifying gene variants or combinations of gene variants that may serve as a screening tool will assist in understanding which individuals will respond as expected to standard c-tDCS protocols and which individuals will not. This may inform the specific c-tDCS parameters to utilise for optimal delivery to all individuals to ensure the number of c-tDCS responders is maximised.

## Author Contributions

Conceived and designed study: MP, SJ, MZ; Carried out data collection: MP; Conducted statistical analysis: MP; Interpreted the findings: MP, SJ, MZ; Wrote the manuscript: MP; Writing and editing of drafts: MP, SJ, MZ.

## References

Abou El Ella, S.S., Tawfik, M.A., Abo El Fotoh, W.M.M., Soliman, O.A.M., 2018. The genetic variant “C588T” of GABARG2 is linked to childhood idiopathic generalized epilepsy and resistance to antiepileptic drugs. Seizure 60, 39–43. https://doi.org/10.1016/j.seizure.2018.06.004

Ammann, C., Lindquist, M.A., Celnik, P.A., 2017. Response variability of different anodal transcranial direct current stimulation intensities across multiple sessions. Brain Stimulation 10, 757–763. https://doi.org/10.1016/j.brs.2017.04.003

Antal, A., Chaieb, L., Moliadze, V., Monte-Silva, K., Poreisz, C., Thirugnanasambandam, N., Nitsche, M.A., Shoukier, M., Ludwig, H., Paulus, W., 2010. Brain-derived neurotrophic factor (BDNF) gene polymorphisms shape cortical plasticity in humans. Brain Stimulation 3, 230–237. https://doi.org/10.1016/j.brs.2009.12.003

Assenza, G., Campana, C., Assenza, F., Pellegrino, G., Di Pino, G., Fabrizio, E., Fini, R., Tombini, M., Di Lazzaro, V., 2017. Cathodal transcranial direct current stimulation reduces seizure frequency in adults with drug-resistant temporal lobe epilepsy: A sham controlled study. Brain Stimulation 10, 333–335. https://doi.org/10.1016/j.brs.2016.12.005

Baghel, R., Grover, S., Kaur, H., Jajodia, A., Rawat, C., Srivastava, A., Kushwaha, S., Agarwal, R., Sharma, S., Kukreti, R., 2016. Evaluating the Role of Genetic Variants on first-line antiepileptic drug response in North India: Significance of *SCN1A* and *GABRA1* Gene Variants in Phenytoin Monotherapy and its Serum Drug Levels. CNS Neurosci Ther 22, 740–757. https://doi.org/10.1111/cns.12570

Bashir, S., Ahmad, S., Alatefi, M., Hamza, A., Sharaf, M., Fecteau, S., Yoo, W.K., 2019. Effects of anodal transcranial direct current stimulation on motor evoked potentials variability in humans. Physiol Rep 7. https://doi.org/10.14814/phy2.14087

Bastani, A., Jaberzadeh, S., 2013. a-tDCS Differential Modulation of Corticospinal Excitability: The Effects of Electrode Size. Brain Stimulation 6, 932–937. https://doi.org/10.1016/j.brs.2013.04.005

Bastani, Andisheh, Jaberzadeh, S., 2013. Differential Modulation of Corticospinal Excitability by Different Current Densities of Anodal Transcranial Direct Current Stimulation. PLoS ONE 8, e72254. https://doi.org/10.1371/journal.pone.0072254

Batsikadze, G., Moliadze, V., Paulus, W., Kuo, M.-F., Nitsche, M.A., 2013. Partially non-linear stimulation intensity-dependent effects of direct current stimulation on motor cortex excitability in humans: Effect of tDCS on cortical excitability. The Journal of Physiology 591, 1987–2000. https://doi.org/10.1113/jphysiol.2012.249730

Brunoni, A.R., Kemp, A.H., Shiozawa, P., Cordeiro, Q., Valiengo, L.C.L., Goulart, A.C., Coprerski, B., Lotufo, P.A., Brunoni, D., Perez, A.B.A., Fregni, F., Benseñor, I.M., 2013. Impact of 5-HTTLPR and BDNF polymorphisms on response to sertraline versus transcranial direct current stimulation: Implications for the serotonergic system. European Neuropsychopharmacology 23, 1530–1540. https://doi.org/10.1016/j.euroneuro.2013.03.009

Cerqueira, V., de Mendonça, A., Minez, A., Dias, A.R., de Carvalho, M., 2006. Does caffeine modify corticomotor excitability? Neurophysiologie Clinique/Clinical Neurophysiology 36, 219–226. https://doi.org/10.1016/j.neucli.2006.08.005

Cheeran, B., Talelli, P., Mori, F., Koch, G., Suppa, A., Edwards, M., Houlden, H., Bhatia, K., Greenwood, R., Rothwell, J.C., 2008. A common polymorphism in the brain-derived neurotrophic factor gene (*BDNF*) modulates human cortical plasticity and the response to rTMS: *BNDF* polymorphism modulates response to rTMS. The Journal of Physiology 586, 5717–5725. https://doi.org/10.1113/jphysiol.2008.159905

Chew, T., Ho, K.-A., Loo, C.K., 2015. Inter- and Intra-individual Variability in Response to Transcranial Direct Current Stimulation (tDCS) at Varying Current Intensities. Brain Stimulation 8, 1130–1137. https://doi.org/10.1016/j.brs.2015.07.031

Chhabra, H., Shivakumar, V., Agarwal, S.M., Bose, A., Venugopal, D., Rajasekaran, A., Subbanna, M., Kalmady, S.V., Narayanaswamy, J.C., Debnath, M., Venkatasubramanian, G., 2016. Transcranial direct current stimulation and neuroplasticity genes: implications for psychiatric disorders. Acta Neuropsychiatr. 28, 1–10. https://doi.org/10.1017/neu.2015.20

Concerto, C., Infortuna, C., Chusid, E., Coira, D., Babayev, J., Metwaly, R., Naenifard, H., Aguglia, E., Battaglia, F., 2017. Caffeinated energy drink intake modulates motor circuits at rest, before and after a movement. Physiology & Behavior 179, 361–368. https://doi.org/10.1016/j.physbeh.2017.07.013

Devanne, H., Cassim, F., Ethier, C., Brizzi, L., Thevenon, A., Capaday, C., 2006. The comparable size and overlapping nature of upper limb distal and proximal muscle representations in the human motor cortex. European Journal of Neuroscience 23, 2467–2476. https://doi.org/10.1111/j.1460-9568.2006.04760.x

Di Lazzaro, V., Manganelli, F., Dileone, M., Notturno, F., Esposito, M., Capasso, M., Dubbioso, R., Pace, M., Ranieri, F., Minicuci, G., Santoro, L., Uncini, A., 2012. The effects of prolonged cathodal direct current stimulation on the excitatory and inhibitory circuits of the ipsilateral and contralateral motor cortex. J Neural Transm 119, 1499–1506. https://doi.org/10.1007/s00702-012-0845-4

Di Lazzaro, V., Pellegrino, G., Di Pino, G., Corbetto, M., Ranieri, F., Brunelli, N., Paolucci, M., Bucossi, S., Ventriglia, M.C., Brown, P., Capone, F., 2015. Val66Met BDNF Gene Polymorphism Influences Human Motor Cortex Plasticity in Acute Stroke. Brain Stimulation 8, 92–96. https://doi.org/10.1016/j.brs.2014.08.006

Di Pino, G., Pellegrino, G., Assenza, G., Capone, F., Ferreri, F., Formica, D., Ranieri, F., Tombini, M., Ziemann, U., Rothwell, J.C., Di Lazzaro, V., 2014. Modulation of brain plasticity in stroke: a novel model for neurorehabilitation. Nat Rev Neurol 10, 597–608. https://doi.org/10.1038/nrneurol.2014.162

Frazer, A., Williams, J., Spittles, M., Rantalainen, T., Kidgell, D., 2016. Anodal transcranial direct current stimulation of the motor cortex increases cortical voluntary activation and neural plasticity: tDCS and Cortical Voluntary Activation. Muscle Nerve 54, 903–913. https://doi.org/10.1002/mus.25143

Fujiyama, H., Hinder, M.R., Barzideh, A., Van de Vijver, C., Badache, A.C., Manrique-C, M.N., Reissig, P., Zhang, X., Levin, O., Summers, J.J., Swinnen, S.P., 2017. Preconditioning tDCS facilitates subsequent tDCS effect on skill acquisition in older adults. Neurobiology of Aging 51, 31–42. https://doi.org/10.1016/j.neurobiolaging.2016.11.012

Fujiyama, H., Hyde, J., Hinder, M.R., Kim, S.-J., McCormack, G.H., Vickers, J.C., Summers, J.J., 2014. Delayed plastic responses to anodal tDCS in older adults. Front. Aging Neurosci. 6. https://doi.org/10.3389/fnagi.2014.00115

Ganho-Ávila, A., Gonçalves, Ó.F., Guiomar, R., Boggio, P.S., Asthana, M.K., Krypotos, A.-M., Almeida, J., 2019. The effect of cathodal tDCS on fear extinction: A cross-measures study. PLoS ONE 14, e0221282. https://doi.org/10.1371/journal.pone.0221282

Gilmore, K.L., Meyers, J.E., 1983. Using Surface Electromyography in Physiotherapy Research. Australian Journal of Physiotherapy 29, 3–9. https://doi.org/10.1016/S0004-9514(14)60659-0

Hair, J., Anderson, R., Tatham, R., Black, W., 1998. Multivariate Data Analysis, 5th ed. Prentice Hall International, Upper Saddle River, NJ, USA.

Hermsen, A.M., Haag, A., Duddek, C., Balkenhol, K., Bugiel, H., Bauer, S., Mylius, V., Menzler, K., Rosenow, F., 2016. Test–retest reliability of single and paired pulse transcranial magnetic stimulation parameters in healthy subjects. Journal of the Neurological Sciences 362, 209–216. https://doi.org/10.1016/j.jns.2016.01.039

Horvath, J.C., Carter, O., Forte, J.D., 2014. Transcranial direct current stimulation: five important issues we aren’t discussing (but probably should be). Front. Syst. Neurosci. 8. https://doi.org/10.3389/fnsys.2014.00002

Hung, C.-C., Chen, P.-L., Huang, W.-M., Tai, J.J., Hsieh, T.-J., Ding, S.-T., Hsieh, Y.-W., Liou, H.-H., 2013. Gene-wide tagging study of the effects of common genetic polymorphisms in the α subunits of the GABA a receptor on epilepsy treatment response. Pharmacogenomics 14, 1849–1856. https://doi.org/10.2217/pgs.13.158

Hwang, J.M., Kim, Y.-H., Yoon, K.J., Uhm, K.E., Chang, W.H., 2015. Different responses to facilitatory rTMS according to BDNF genotype. Clinical Neurophysiology 126, 1348–1353. https://doi.org/10.1016/j.clinph.2014.09.028

Inghilleri, M., Conte, A., Currà, A., Frasca, V., Lorenzano, C., Berardelli, A., 2004. Ovarian hormones and cortical excitability. An rTMS study in humans. Clinical Neurophysiology 115, 1063–1068. https://doi.org/10.1016/j.clinph.2003.12.003

Keel, J., Smith, M., Wassermann, E., 2001. A safety screening questionnaire for transcranial magnetic stimulation. Clinical Neurophysiology 112, 720. https://doi.org/10.1016/S1388-2457(00)00518-6

Kendell, F., McCreary, E., Provance, P., 2010. Muscles, testing and function with posture and pain. Williams and Wilkins, Baltimore, MD, USA.

Kujirai, T., Caramia, M.D., Rothwell, J.C., Day, B.L., Thompson, P.D., Ferbert, A., Wroe, S., Asselman, P., Marsden, C.D., 1993. Corticocortical inhibition in human motor cortex. The Journal of Physiology 471, 501–519. https://doi.org/10.1113/jphysiol.1993.sp019912

Labruna, L., Jamil, A., Fresnoza, S., Batsikadze, G., Kuo, M.-F., Vanderschelden, B., Ivry, R.B., Nitsche, M.A., 2016. Efficacy of Anodal Transcranial Direct Current Stimulation is Related to Sensitivity to Transcranial Magnetic Stimulation. Brain Stimulation 9, 8–15. https://doi.org/10.1016/j.brs.2015.08.014

Lee, L.-C., Cho, Y.-C., Lin, P.-J., Yeh, T.-C., Chang, C.-Y., Yeh, T.-K., 2016. Influence of Genetic Variants of the N-Methyl-D-Aspartate Receptor on Emotion and Social Behavior in Adolescents. Neural Plasticity 2016, 1–8. https://doi.org/10.1155/2016/6851592

Li, L.M., Uehara, K., Hanakawa, T., 2015. The contribution of interindividual factors to variability of response in transcranial direct current stimulation studies. Front. Cell. Neurosci. 9. https://doi.org/10.3389/fncel.2015.00181

Lin, L.-C., Ouyang, C.-S., Chiang, C.-T., Yang, R.-C., Wu, R.-C., Wu, H.-C., 2018. Cumulative effect of transcranial direct current stimulation in patients with partial refractory epilepsy and its association with phase lag index-A preliminary study. Epilepsy & Behavior 84, 142–147. https://doi.org/10.1016/j.yebeh.2018.04.017

López-Alonso, V., Cheeran, B., Río-Rodríguez, D., Fernández-del-Olmo, M., 2014. Inter-individual Variability in Response to Non-invasive Brain Stimulation Paradigms. Brain Stimulation 7, 372–380. https://doi.org/10.1016/j.brs.2014.02.004

López-Alonso, V., Fernández-del-Olmo, M., Costantini, A., Gonzalez-Henriquez, J.J., Cheeran, B., 2015. Intra-individual variability in the response to anodal transcranial direct current stimulation. Clinical Neurophysiology 126, 2342–2347. https://doi.org/10.1016/j.clinph.2015.03.022

Mastroeni, C., Bergmann, T.O., Rizzo, V., Ritter, C., Klein, C., Pohlmann, I., Brueggemann, N., Quartarone, A., Siebner, H.R., 2013. Brain-Derived Neurotrophic Factor – A Major Player in Stimulation-Induced Homeostatic Metaplasticity of Human Motor Cortex? PLoS ONE 8, e57957. https://doi.org/10.1371/journal.pone.0057957

Matamala, J.M., Howells, J., Dharmadasa, T., Trinh, T., Ma, Y., Lera, L., Vucic, S., Burke, D., Kiernan, M.C., 2018. Inter-session reliability of short-interval intracortical inhibition measured by threshold tracking TMS. Neuroscience Letters 674, 18–23. https://doi.org/10.1016/j.neulet.2018.02.065

McDonnell, M.N., Orekhov, Y., Ziemann, U., 2006. The role of GABAB receptors in intracortical inhibition in the human motor cortex. Exp Brain Res 173, 86–93. https://doi.org/10.1007/s00221-006-0365-2

Monte-Silva, K., Kuo, M.-F., Hessenthaler, S., Fresnoza, S., Liebetanz, D., Paulus, W., Nitsche, M.A., 2013. Induction of Late LTP-Like Plasticity in the Human Motor Cortex by Repeated Non-Invasive Brain Stimulation. Brain Stimulation 6, 424–432. https://doi.org/10.1016/j.brs.2012.04.011

Mori, F., Ribolsi, M., Kusayanagi, H., Siracusano, A., Mantovani, V., Marasco, E., Bernardi, G., Centonze, D., 2011. Genetic variants of the NMDA receptor influence cortical excitability and plasticity in humans. Journal of Neurophysiology 106, 1637–1643. https://doi.org/10.1152/jn.00318.2011

Naimo, G.D., Guarnaccia, M., Sprovieri, T., Ungaro, C., Conforti, F.L., Andò, S., Cavallaro, S., 2019. A Systems Biology Approach for Personalized Medicine in Refractory Epilepsy. IJMS 20, 3717. https://doi.org/10.3390/ijms20153717

Narita, S., Onozawa, Y., Yoshihara, E., Nishizawa, D., Numajiri, M., Ikeda, K., Iwahashi, K., 2018. Association between N-methyl-d-aspartate Receptor Subunit 2B Gene Polymorphisms and Personality Traits in a Young Japanese Population. East Asian Arch Psychiatry 28, 45–52. https://doi.org/10.12809/eaap181712

Nitsche, M.A., Cohen, L.G., Wassermann, E.M., Priori, A., Lang, N., Antal, A., Paulus, W., Hummel, F., Boggio, P.S., Fregni, F., Pascual-Leone, A., 2008. Transcranial direct current stimulation: State of the art 2008. Brain Stimulation 1, 206–223. https://doi.org/10.1016/j.brs.2008.06.004

Nitsche, M.A., Doemkes, S., Karaköse, T., Antal, A., Liebetanz, D., Lang, N., Tergau, F., Paulus, W., 2007. Shaping the Effects of Transcranial Direct Current Stimulation of the Human Motor Cortex. Journal of Neurophysiology 97, 3109–3117. https://doi.org/10.1152/jn.01312.2006

Nitsche, M.A., Fricke, K., Henschke, U., Schlitterlau, A., Liebetanz, D., Lang, N., Henning, S., Tergau, F., Paulus, W., 2003. Pharmacological Modulation of Cortical Excitability Shifts Induced by Transcranial Direct Current Stimulation in Humans. The Journal of Physiology 553, 293–301. https://doi.org/10.1113/jphysiol.2003.049916

Nitsche, M.A., Paulus, W., 2000. Excitability changes induced in the human motor cortex by weak transcranial direct current stimulation. The Journal of Physiology 527, 633–639. https://doi.org/10.1111/j.1469-7793.2000.t01-1-00633.x

Oldfield, R., 1971. The assessment and analysis of handedness: The Ediburgh inventory 9, 97–113.

O’leary, T.J., Morris, M.G., Collett, J., Howells, K., 2015. Reliability of single and paired-pulse transcranial magnetic stimulation in the vastus lateralis muscle: Intracortical Inhibition and Facilitation. Muscle & Nerve 52, 605–615. https://doi.org/10.1002/mus.24584

Pellegrini, M., Zoghi, M., Jaberzadeh, S., 2020a. Can genetic polymorphisms predict response variability to anodal transcranial direct current stimulation of the primary motor cortex? (preprint). Neuroscience. https://doi.org/10.1101/2020.03.31.017798

Pellegrini, M., Zoghi, M., Jaberzadeh, S., 2020b. The effects of transcranial direct current stimulation on corticospinal and cortico-cortical excitability and response variability: conventional versus high definition montages (preprint). Neuroscience. https://doi.org/10.1101/2020.03.30.017046

Pellegrini, M., Zoghi, M., Jaberzadeh, S., 2020c. A Checklist to Reduce Response Variability in Studies Using Transcranial Magnetic Stimulation for Assessment of Corticospinal Excitability: A Systematic Review of the Literature. Brain Connectivity 10, 53–71. https://doi.org/10.1089/brain.2019.0715

Pellegrini, M., Zoghi, M., Jaberzadeh, S., 2018a. Biological and anatomical factors influencing interindividual variability to noninvasive brain stimulation of the primary motor cortex: a systematic review and meta-analysis. Reviews in the Neurosciences 29, 199–222. https://doi.org/10.1515/revneuro-2017-0048

Pellegrini, M., Zoghi, M., Jaberzadeh, S., 2018b. Cluster analysis and subgrouping to investigate inter-individual variability to non-invasive brain stimulation: a systematic review. Reviews in the Neurosciences 29, 675–697. https://doi.org/10.1515/revneuro-2017-0083

Pellegrini, M., Zoghi, M., Jaberzadeh, S., 2018c. The effect of transcranial magnetic stimulation test intensity on the amplitude, variability and reliability of motor evoked potentials. Brain Research 1700, 190–198. https://doi.org/10.1016/j.brainres.2018.09.002

Portney, L., Watkins, M., 2009. Foundations of Clinical Research: applications to Practice., 3rd ed. Prentice Hall, Upper Saddle River, NJ, USA.

Puri, R., Hinder, M.R., Canty, A.J., Summers, J.J., 2016. Facilitatory non-invasive brain stimulation in older adults: the effect of stimulation type and duration on the induction of motor cortex plasticity. Experimental Brain Research 234, 3411–3423. https://doi.org/10.1007/s00221-016-4740-3

Puri, R., Hinder, M.R., Fujiyama, H., Gomez, R., Carson, R.G., Summers, J.J., 2015. Duration-dependent effects of the BDNF Val66Met polymorphism on anodal tDCS induced motor cortex plasticity in older adults: a group and individual perspective. Frontiers in Aging Neuroscience 7. https://doi.org/10.3389/fnagi.2015.00107

Ridding, M.C., Ziemann, U., 2010. Determinants of the induction of cortical plasticity by non-invasive brain stimulation in healthy subjects: Induction of cortical plasticity by non-invasive brain stimulation. The Journal of Physiology 588, 2291–2304. https://doi.org/10.1113/jphysiol.2010.190314

Rossi, S., Studer, V., Moscatelli, A., Motta, C., Coghe, G., Fenu, G., Caillier, S., Buttari, F., Mori, F., Barbieri, F., Castelli, M., De Chiara, V., Monteleone, F., Mancino, R., Bernardi, G., Baranzini, S.E., Marrosu, M.G., Oksenberg, J.R., Centonze, D., 2013. Opposite Roles of NMDA Receptors in Relapsing and Primary Progressive Multiple Sclerosis. PLoS ONE 8, e67357. https://doi.org/10.1371/journal.pone.0067357

Rossini, P.M., Rossi, S., 1998. Clinical applications of motor evoked potentials. Electroencephalography and Clinical Neurophysiology 106, 180–194. https://doi.org/10.1016/S0013-4694(97)00097-7

Rothwell, J., Hallett, M., Berardelli, A., Eisen, A., Rossini, P., Paulus, W., 1999. Magnetic stimulation: motor evoked potentials, in: Recommendations for the Practice of Clinical Neurophysiology: Guidelines of the International Federation of Clinical Neurophysiology.

Sale, M.V., Ridding, M.C., Nordstrom, M.A., 2008. Cortisol Inhibits Neuroplasticity Induction in Human Motor Cortex. Journal of Neuroscience 28, 8285–8293. https://doi.org/10.1523/JNEUROSCI.1963-08.2008

Sale, M.V., Ridding, M.C., Nordstrom, M.A., 2007. Factors influencing the magnitude and reproducibility of corticomotor excitability changes induced by paired associative stimulation. Experimental Brain Research 181, 615–626. https://doi.org/10.1007/s00221-007-0960-x

Sasaki, R., Miyaguchi, S., Kotan, S., Kojima, S., Kirimoto, H., Onishi, H., 2016. Modulation of Cortical Inhibitory Circuits after Cathodal Transcranial Direct Current Stimulation over the Primary Motor Cortex. Front. Hum. Neurosci. 10. https://doi.org/10.3389/fnhum.2016.00030

Smith, M.J., Adams, L.F., Schmidt, P.J., Rubinow, D.R., Wassermann, E.M., 2002. Effects of ovarian hormones on human cortical excitability. Annals of Neurology 51, 599–603. https://doi.org/10.1002/ana.10180

SPSS, 2001. The SPSS TwoStep Cluster Component: A scalble component enabling more efficient customer segmentation. SPSS Inc.

Strube, W., Bunse, T., Malchow, B., Hasan, A., 2015. Efficacy and Interindividual Variability in Motor-Cortex Plasticity following Anodal tDCS and Paired-Associative Stimulation. Neural Plasticity 2015, 1–10. https://doi.org/10.1155/2015/530423

Strube, W., Bunse, T., Nitsche, M.A., Nikolaeva, A., Palm, U., Padberg, F., Falkai, P., Hasan, A., 2016. Bidirectional variability in motor cortex excitability modulation following 1 mA transcranial direct current stimulation in healthy participants. Physiological Reports 4, e12884. https://doi.org/10.14814/phy2.12884

Tecchio, F., Cottone, C., Porcaro, C., Cancelli, A., Di Lazzaro, V., Assenza, G., 2018. Brain Functional Connectivity Changes After Transcranial Direct Current Stimulation in Epileptic Patients. Front. Neural Circuits 12, 44. https://doi.org/10.3389/fncir.2018.00044

Tekturk, P., Erdogan, E.T., Kurt, A., Kocagoncu, E., Kucuk, Z., Kinay, D., Yapici, Z., Aksu, S., Baykan, B., Karamursel, S., 2016. Transcranial direct current stimulation improves seizure control in patients with Rasmussen encephalitis. Epileptic Disorders 18, 58–66. https://doi.org/10.1684/epd.2016.0796

Teo, J.T.H., Bentley, G., Lawrence, P., Soltesz, F., Miller, S., Willé, D., McHugh, S., Dodds, C., Lu, B., Croft, R.J., Bullmore, E.T., Nathan, P.J., 2014. Late cortical plasticity in motor and auditory cortex: role of met-allele in BDNF Val66Met polymorphism. Int. J. Neuropsychopharm. 17, 705–713. https://doi.org/10.1017/S1461145713001636

Tremblay, S., Larochelle-Brunet, F., Lafleur, L.-P., El Mouderrib, S., Lepage, J.-F., Théoret, H., 2016. Systematic assessment of duration and intensity of anodal transcranial direct current stimulation on primary motor cortex excitability. European Journal of Neuroscience 44, 2184–2190. https://doi.org/10.1111/ejn.13321

Vaseghi, B., Zoghi, M., Jaberzadeh, S., 2016. Unihemispheric concurrent dual-site cathodal transcranial direct current stimulation: the effects on corticospinal excitability. Eur J Neurosci 43, 1161–1172. https://doi.org/10.1111/ejn.13229

Wiethoff, S., Hamada, M., Rothwell, J.C., 2014. Variability in Response to Transcranial Direct Current Stimulation of the Motor Cortex. Brain Stimulation 7, 468–475. https://doi.org/10.1016/j.brs.2014.02.003

Yook, S.-W., Park, S.-H., Seo, J.-H., Kim, S.-J., Ko, M.-H., 2011. Suppression of Seizure by Cathodal Transcranial Direct Current Stimulation in an Epileptic Patient - A Case Report -. Ann Rehabil Med 35, 579. https://doi.org/10.5535/arm.2011.35.4.579

Zoghi, M., O’Brien, T.J., Kwan, P., Cook, M.J., Galea, M., Jaberzadeh, S., 2016. Cathodal transcranial direct-current stimulation for treatment of drug-resistant temporal lobe epilepsy: A pilot randomized controlled trial. Epilepsia Open 1, 130–135. https://doi.org/10.1002/epi4.12020

Zoghi, M., Vaseghi, B., Bastani, A., Jaberzadeh, S., Galea, M.P., 2015. The Effects of Sex Hormonal Fluctuations during Menstrual Cycle on Cortical Excitability and Manual Dexterity (a Pilot Study). PLOS ONE 10, e0136081. https://doi.org/10.1371/journal.pone.0136081

